# Structural basis of the protochromic green/red photocycle of the chromatic acclimation sensor RcaE

**DOI:** 10.1101/2020.12.08.414581

**Authors:** Takayuki Nagae, Masashi Unno, Taiki Koizumi, Yohei Miyanoiri, Tomotsumi Fujisawa, Kento Masui, Takanari Kamo, Kei Wada, Toshihiko Eki, Yutaka Ito, Yuu Hirose, Masaki Mishima

**Affiliations:** Synchrotron Radiation Research Center, Nagoya University, Chikusa, Nagoya 464-8603, Japan; Department of Chemistry and Applied Chemistry, Faculty of Science and Engineering, Saga University, Saga 840-8502, Japan; Department of Chemistry, Graduate School of Science, Tokyo Metropolitan University, 1-1 Minamiosawa, Hachioji 192-0397, Japan; Institute for Protein Research, Osaka University, 3-2 Yamadaoka, Suita, Osaka 565-0871, Japan; Department of Medical Sciences, University of Miyazaki, Miyazaki 889-1692, Japan; Department of Environmental and Life Sciences, Toyohashi University of Technology, Toyohashi, Aichi 441-8580, Japan

**Keywords:** cyanobacteriochrome, phytochrome, bilin, NMR, crystallography

## Abstract

Cyanobacteriochromes (CBCRs) are bilin-binding photosensors of the phytochrome superfamily that show remarkable spectral diversity. The green/red CBCR subfamily is important for regulating chromatic acclimation of photosynthetic antenna in cyanobacteria and is applied for optogenetic control of gene expression in synthetic biology. They are suggested to combine the bilin C15-*Z*/C15-*E* photoisomerization with a change in the bilin protonation state to drive their absorption changes. However, structural information and direct evidence of the bilin protonation state are lacking. Here we report a high-resolution (1.63Å) crystal structure of the bilin-binding domain of the chromatic acclimation sensor RcaE in the red-absorbing photoproduct state. The bilin is buried within a “pan” consisting of hydrophobic residues, where the bilin configuration/conformation is C5-*Z,syn*/C10-*Z,syn/*C15-*E,syn* with the A–C rings co-planar and the D-ring tilted. Three pyrrole nitrogens of the A–C rings are covered in the α-face with a hydrophobic lid of Leu249 influencing the bilin p*K*_a_, whereas they are directly hydrogen-bonded in the β-face with the carboxyl group of Glu217. Glu217 is further connected to a cluster of waters forming a hole in the pan, which are in exchange with solvent waters in molecular dynamics simulation. We propose that the “holey pan” structure functions as a proton-exit/influx pathway upon photoconversion. NMR analysis demonstrated that the four pyrrole nitrogen atoms are indeed fully protonated in the red-absorbing state, but one of them, most likely the B-ring nitrogen, is deprotonated in the green-absorbing state. These findings deepen our understanding of the diverse spectral tuning mechanisms present in CBCRs.

**Significance Statement:** Green/red CBCRs are one of the most important CBCR subfamilies owing to their physiological roles in cyanobacteria phylum and optogenetic applications. They are known to utilize a change in the bilin protonation state to drive the marked change in green/red absorption, but the structural basis of the protochromic green/red photocycle are not well understood. Here, we have determined the crystal structure of the chromatic acclimation sensor RcaE of this subfamily in the photoproduct state, demonstrating a unique conformation of the bilin and its interacting residues. In addition, we provide direct evidence of the protonation state of the bilin via NMR analysis. These findings bring insight to our understanding of the molecular mechanisms underlying the spectral diversity of CBCRs.

## Introduction

Phytochromes and cyanobacteriochromes (CBCRs) are members of phytochrome superfamily of photosensors. They utilize a cysteine-linked linear tetrapyrrole (bilin) chromophore for photo-perception that is bound within a GAF (cGMP phosphodiesterase/adenylyl cyclase/FhlA) domain (1–6). They undergo reversible photoconversion between two distinct light-absorbing states triggered by C15*-Z*/C15*-E* photoisomerization of the bilin chromophore (7, 8). Phytochromes are distributed in higher plants, algae, bacteria, and fungi; they typically photoconvert between a red-absorbing C15*-Z,anti* dark state and a far-red light-absorbing C15*-E,anti* photoproduct state, although recent studies have revealed substantial spectral diversity in some algal and cyanobacterial phytochromes (9, 10). CBCRs are widely distributed among the cyanobacteria phylum, and show marked variation in their absorbing wavelength, spanning the near-UV to the far-red part of the spectrum (reviewed recently in (4–6)). They require only a GAF domain to complete the photocycle, in contrast to phytochromes, which require an extra PHY (phytochrome) domain and, in many cases, a PAS (Per/Arnt/Sim) domain to complete their photocycle (11, 12). The GAF domains of CBCRs have been classified into subfamilies based on the configurations of their dark- and photoproduct-state chromophores, designated by their respective light-absorbing maxima (e.g., blue/green, green/red, etc. subfamilies).

To date, several chemical reactions are known to create the spectral diversity of the CBCR subfamilies, most of which use the chromophore precursor, phycocyanobilin. The “two-Cys photocycle” mechanism was first proposed in the blue/green subfamily by Rockwell et al. (13) and this mechanism appeared widespread among other violet- and blue-absorbing CBCR subfamilies (14). As confirmed by structural studies, the thiol group of a second Cys residue forms a thioether linkage with the C10 atom, disconnecting bilin conjugation between the B- and C-rings to cause a blue shift in absorption (15, 16). Some CBCR members undergoing two-Cys photocycle isomerize their PCB chromophore to phycoviolobilin (17), which disconnects the π-conjugated system between the A- and B-rings and leads to spectral blue-shift in their Cys-free photostates. Subsequently, the “protochromic photocycle” was identified in the green/red subfamily in our previous study (18). In this mechanism, a change in the protonation state of the bilin pyrrole system is responsible for the change in absorption. The deprotonation-induced formation of green-absorbing state has also been observed as an intermediate in the two-Cys photocycle (19). The red/green subfamily harbors a fully protonated chromophore in both photostates, but a twisted conformation of the A- and D-rings is responsible for the blue-shifted photoproduct (8, 20–23); this is known as the “trapped-twist photocycle” (24). Recently, Bandara et al. reported the structure of a far-red absorbing state of a CBCR that adopts a atypical C15*-Z*,*syn* configuration (25). They proposed a model that its far-red absorption maximum arises from a cationic bilin tautomer in which the A-ring is formally di-protonated while the B-ring remains deprotonated.

Cyanobacteria harbor up to several dozen different bilin-binding GAF domains (26), and a single CBCR protein often contains multiple GAF domains (27), suggesting that cyanobacteria utilize a highly complex photosensing system. Although some CBCRs have been shown to regulate the composition of photosynthetic antenna (4, 28–31), cell morphology (32), phototaxis (33), and cell aggregation (34, 35), the physiological role of most CBCRs remains unknown. RcaE and CcaS are members of the green/red CBCR subfamily and optimize the light-absorption maxima of the photosynthetic antenna complex phycobilisome via transcriptional regulation of its components in a process called “chromatic acclimation”. RcaE controls the expression of phycobilisome genes via phosphorylation of RcaF and RcaC under red light (4, 18, 27, 28, 36, 37), whereas CcaS controls fewer genes via phosphorylation of the transcriptional factor CcaR under green light (38, 39). The CcaSR system has been applied as one of the most popular photoswitch systems of gene expression in synthetic biology (40–42). RcaE and/or CcaS orthologs are found in ~15% of all sequenced cyanobacteria genomes, suggesting that they have a physiological impact on survival (43).

Based on pH titration experiments, we previously demonstrated that the bilin-binding GAF domain of RcaE exhibits a protochromic photocycle consisting of a deprotonated C15*-Z* Pg dark state and a protonated C15*-E* Pr photoproduct state(18). In this photocycle, green or red absorption is determined by the protonation state of bilin, with C15*-Z*/C15*-E* photoisomerization responsible for tuning its p*K*_a_(18). Site-directed mutagenesis of RcaE identified the protochromic triad residues, Glu217, Leu249 and K261, required for the marked shift in the bilin p*K*_a_(18). Resonance Raman spectroscopy of the C15*-Z* Pg dark state of RcaE (44) was highly consistent with B-ring-deprotonated bilin system that adopts a C15*-Z,anti* structure based upon QM/MM calculations using the structure of C15*-Z*,*anti* Pr dark state of AnPixJ as template (45). However, structural information and direct evidence of the chemical environments of bilin in the green/red CBCR subfamily have not yet been reported, preventing deeper understanding of the molecular basis of the protochromic photocycle.

In this study, we report the crystal structure of the RcaE GAF domain in its Pr photoproduct state, in which its PCB chromophore adopts the C15*-E*,*syn* structure never seen previously. We also discovered that PCB is connected to a unique water cluster via hydrogen bond network via Glu217, which may function as a proton-exit/influx pathway upon photoconversion. Nuclear magnetic resonance (NMR) analysis demonstrated that all four pyrrole nitrogen atoms of the bilin are indeed fully protonated in the Pr photoproduct state, whereas one is deprotonated in the Pg state. These findings provide new insight into our understanding of the diverse spectral tuning mechanisms in CBCRs.

## Results

### Crystal Structure of the GAF Domain

The GAF domain of RcaE was recombinantly expressed in PCB-producing *E. coli* and purified by Ni-affinity and gel filtration chromatography (46) (Fig. 1A). The protein was crystallized in the Pr photoproduct state and X-ray diffraction data were collected (see Materials and Methods for detail). The initial phase was derived by molecular replacement using the coordinates of the TePixJ GAF domain (PDB: 3VV4) as a search model, and the model was refined to a resolution of 1.63 Å and an *R* value of 15.8% (free *R*, 19.9%) (Table S1). Two protomers whose structures were essentially identical (r.m.s.d. for 139 Cα atoms, 0.17 Å) were contained in the asymmetric unit, and molecule A is described below as a representative. Molecule A includes residues 164–268 and 277–313, and covalently bound PCB. The mainframe of the structure displays a typical GAF fold composed of a five-stranded antiparallel β-sheet and five helices (Fig. 1B and C). Among the five helices, the short second helix, named H2, is a single turn of a 3_10_ helix (Fig. 1B and C). The five β-sheet forms aromatic and aliphatic hydrophobic cores on both sides, and is backed by the H1 and H5 helices (Fig. 1B). PCB is covalently anchored at a cysteine residue in the H4 helix and buried within the cleft in the GAF domain. The α-face of PCB is covered by residues from the H4 helix (Fig. 1B and D). The β-face of PCB is covered by a core of the antiparallel β-sheet, which is constituted by strand 4 (S4), strand 5 (S5), and strand 1 (S1), and also a large loop between the S2-S3 sheets (S2-S3 loop, residues 203–232) that include the H2 and H3 helices (Fig. 1B and D). This structural arrangement of the PCB-binding pocket invokes a “lidded pan”, where the H4 helix forms the lid, and the S4, S5, S1 strands and the S2-S3 loop together form the pan.

**Figure 1.**
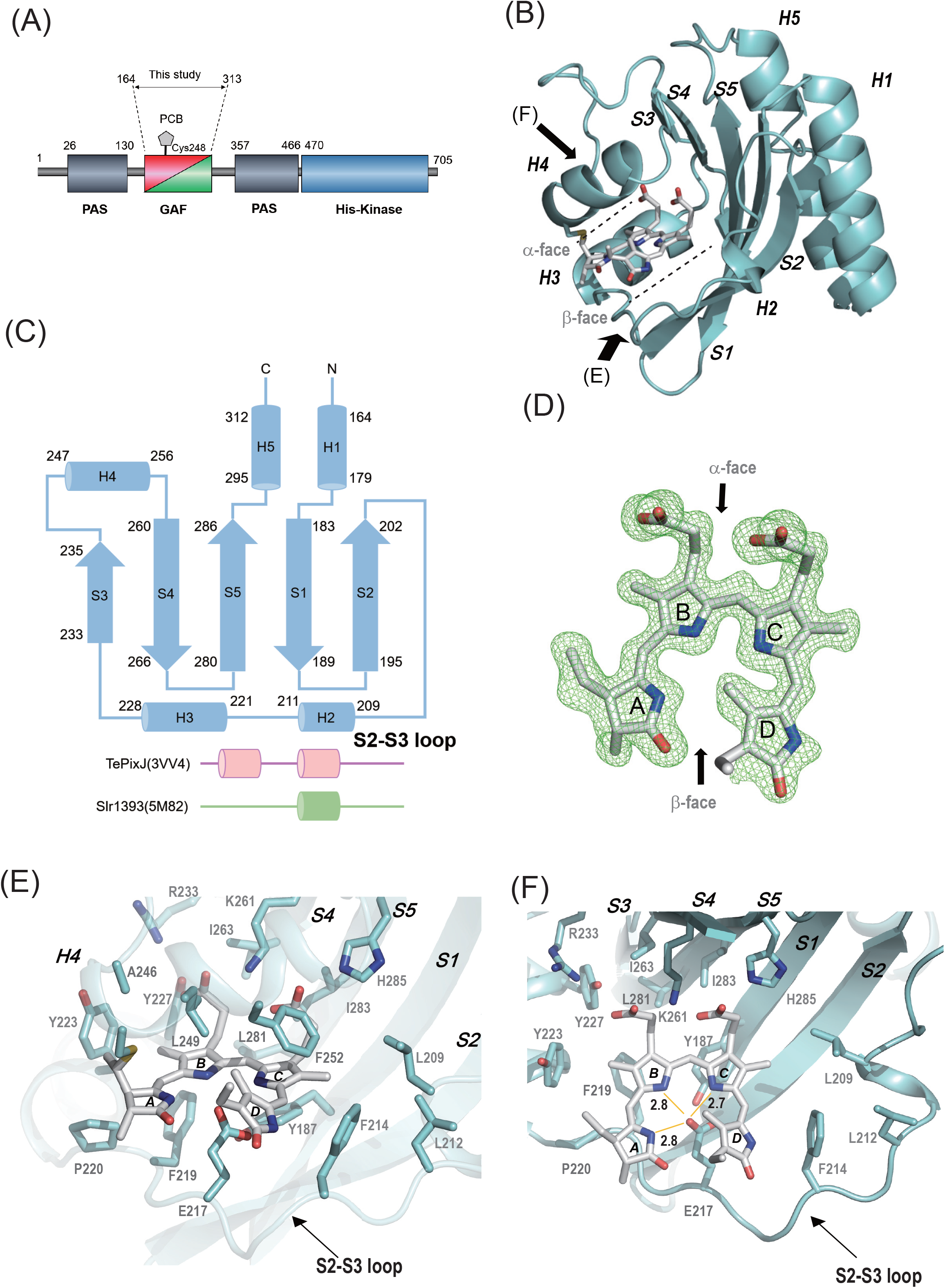
Structure of the chromatic acclimation sensor RcaE. (A) Domain structure of RcaE. (B) Ribbon drawings of the crystal structure of the GAF domain of RcaE (cyan). The bound PCB molecule is depicted in stick representation (gray).The α-face and β-face that account for each side of PCB are indicated by dotted lines. Arrows indicate the direction viewed in panels E and F. (C) Topology diagram of the secondary structure of the GAF domain of RcaE (cyan). The regions of TePixJ and Slr1393 corresponding to the S2-S3 loop of RcaE are shown in pink and green, respectively (D) F_o_-F_c_ map of PCB contoured at 2.5σ. The α-face and β-face are indicated by arrows. (E) Close-up view of the PCB-binding site along the arrow labeled E in panel A. (F) Close-up view shown along the black arrow labeled F in panel A. The distances between the oxygen atom of carboxyl group of the Glu217 sidechain and NA, NB, and NC of the PCB are shown in Å. For clarity, H4 has been omitted from the figures (E) and (F).

### Detailed structure of PCB and its interacting residues

The clear electron density map and F_o_-F_c_ map allowed us to determine the conformations of the PCB chromophore in detail (Fig. 1D). In general, the geometry of the A-to D-rings of the bilin chromophore are each classified by *syn/anti* conformation and *Z/E* configuration at the C5, C10, and C15 atoms (64 possible classifications in total). In the Pr structure of RcaE, PCB adopts C5-*Z*, C10-*Z*, and C15-*E* configurations, but remarkably it adopts all *syn* conformations. To our knowledge, this is the first report of the bilin-binding protein that harbors a C15-*E*,*syn* chromophore. The C15-*E* configuration for the Pr state is consistent with previous results from acid denaturation analysis (18, 38). The A-, B-, and C-rings of PCB form a nearly planar structure, but the D ring is tilted by approximately 23 degrees out of the A-B-C plane (Fig. 1D). The hydrophobic lid covering the α-face of PCB is composed of two residues, Leu249 and Phe252 (Fig. 1E). The tilt of the D-ring is supported by hydrophobic interactions with Phe252 on the H4 helix (Fig. 1E). The methyl group of C17 of the D-ring makes Van der Waals contacts with the methyl group of Leu249. The amide group of the D-ring faces the aromatic ring of Phe214 on the S2-S3 loop (Fig. 1E and F).

The location of PCB against the protein in RcaE is distinct in comparison to other CBCRs. When the structure is viewed with the H4 helix side-up (from the α-face of PCB), PCB is rotated clockwise by one pyrrole ring (Fig. 2A). For structural comparison, we chose the GAF domains of TePixJ (PDB:3VV4) and Slr1393 (PDB:5M82), which harbor a bilin chromophore in the C15-*E* configuration (23, 45). Essentially, the protein part of the GAF domain of RcaE is well fitted to the other GAF domains, but the position of PCB in RcaE is strikingly different. The distances between TePixJ and RcaE with respect to the representative PCB atoms range from 3.1 to 4.1 Å (Table S2). Similarly, the distances between Slr1393 and RcaE with respect to the representative PCB atoms range from 3.2 to 5.1 Å. Recently, Bandara et al. reported the far-red-absorbing state of the 2551g3 of far-red/orange CBCR subfamily, whose GAF domain shows high sequence similarity to that of RcaE. The Pfr dark state of 2551g3 adopts all *syn* conformation and all *Z* configuration, but also shows a clockwise-rotated location of PCB similar to that seen in RcaE (25).

**Figure 2.**
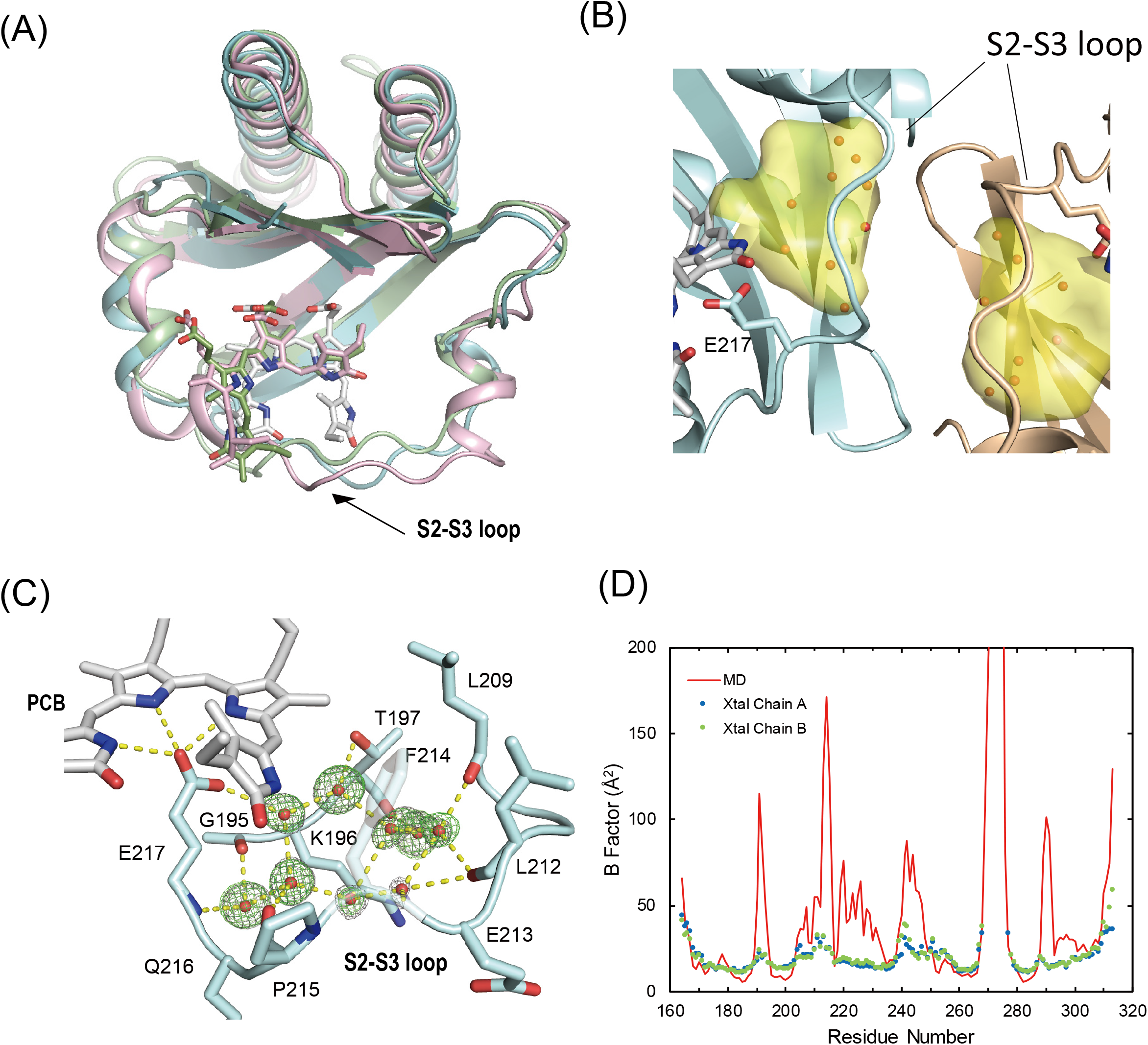
Features of the structure of RcaE. (A) Superimposition of the crystal structure of the GAF domain of RcaE (cyan) on those of TePixJ (green) and Slr1393 (pink). The H4 of RcaE and the corresponding region of TePixJ and Slr1393 have been omitted for clarity. (B) The cavity inside the protein shown as surface representations in yellow. The chain A of the GAF domain is represented as a cyan ribbon. The chain B of a neighboring asymmetric unit is represented as brown. The bound PCB molecules are shown as stick models colored in gray. (C) Clustered waters (oxygen) inside the S2-S3 loop of chain A shown as red balls. Electron-density and difference electron-density map are also shown as gray and green meshes contoured at 1.0σ and 3.0σ, respectively. The hydrogen-bond network is illustrated by yellow dashed lines. (D) Calculated average B-factors per residue. MD values, and values for chain A, and chain B obtained from the crystal structure are shown in red, blue and green, respectively.

The key interactions contributing to the location of PCB in the GAF domain of RcaE are hydrogen bonds and/or ion pairs between the carboxylates of the propionic acid of the B-ring and C-ring (hereafter called B-carboxylate and C-carboxylate, respectively). The Tyr227 sidechain forms a hydrogen bond with the B-carboxylate (Fig. 1F), and the Lys261 sidechain make ion pairs with the B- and C-carboxylates (Fig. 1F). The distance between the Nε atom of the His285 sidechain and C-carboxylate is 2.7 Å, and thus the His285 sidechain probably also makes an ion pair or hydrogen bond with the C-carboxylate (Fig. 1 E and F). Glu217, the sole acidic interactor with PCB, makes hydrogen bonds with PCB: the distances between one oxygen atom of the carboxyl group of the Glu217 sidechain and N_A_ (A-ring), N_B_ (B-ring), and N_C_ (C-ring) are 2.8 Å, 2.8 Å, and 2.7 Å, respectively (Fig. 1F). Of note, the corresponding distance for N_D_ (D-ring) is 4.4 Å. The other oxygen on the opposite side of Glu217 forms a hydrogen bond with Tyr187 in the aromatic ring (Fig. 1F). Furthermore, the aromatic ring of Tyr223 interacts with the methyl group of the B-ring in a perpendicular manner, probably by forming a CH–π interaction (Fig. 1E and F). The sidechains of Leu209 and Phe214 on the S2-S3 loop make the part of hydrophobic pan, which encloses the methyl group of the C-ring and the amide group of the D-ring (Fig. 1F).

### Identification of the “holey pan” structure

The hydrophobic residues on the S2-S3 loop, namely Leu209, Phe214, Phe219, Pro220, and Tyr223, form the wall of the hydrophobic pan, which surrounds PCB without any direct hydrogen bonds (Fig. 1E and F). Because the packing of atoms in the hydrophobic region seems loose, we surveyed the cavity inside the GAF domain by using CAVER (47), a program that can detect interior cavities of proteins. Notably, we identified a cavity between the C- and D-rings, the S2-S3 loop, and the S2 strand, with a size of 196.9 Å^3^ (Fig. 2B, in yellow). This large protein cavity was filled an ordered cluster of water molecules that contains a pentagonal core structure (Fig. 2B and C). Although S2-S3 loop of chain A partly interacts with a neighboring molecule, the water molecules penetrated into the cavity are not directly involved in the crystal-packing contact (Fig. 2B). This “pentagon water” has been seen in other proteins at their hydrophobic surface by X-ray crystallography (48, 49). However, a similar cavity incorporating a water cluster has not been reported in any other structures of phytochromes or CBCRs including the Pfr dark state of 2551g3 (25). The amino acid sequences of the S2-S3 loop, such as the hydrophobic residues Leu209, Phe214, Phe219, Pro220, and Tyr223 are well conserved among green/red CBCRs, but highly diverse among the other CBCRs including the far-red/orange subfamily (Fig. S2). Namely, we identified a “hole” in the hydrophobic pan of the Pr structure of RcaE, which is conserved specifically in the green/red CBCRs.

Next, we investigated the dynamic properties of the GAF domain, especially the key interactions between PCB and the protein, the S2-S3 loop, and the clustered water molecules by molecular dynamics (MD) simulation. A 500-ns MD run at 300 K was performed using the crystal structure as an initial starting geometry and including explicit solvent water molecules. The ion pairs between Lys261 and the B- and C-carboxylates were stable throughout the MD simulation, although the carboxylate rotates to replace each oxygen with the other (Fig. S3A, Supplementary Movie). Furthermore, the sole acidic interactor Glu217 still maintained interactions with PCB, even though the carboxylate oxygen atoms were in exchange (Fig. S3B, Supplementary Movie).

The root mean square fluctuation (rmsf) of the backbone C, N, and O atoms over the whole trajectory was calculated and converted to average B-factors, which were used as a measure of structural flexibility. Nearly all of the residues in the GAF domain had relatively low average B-factors with the exception of residues close to the N and C termini, and residues 269–276, for which the coordinates are missing in the crystal structure (Fig. 2D). Interestingly, the average B-factor was larger for residues Leu209 to Phe219, corresponding to the middle portion of the S2-S3 loop, than for other parts of the domain (Fig. 2D). Notably, the experimentally derived B-factors from the X-ray crystallographic analysis showed a similar pattern to that of the MD simulation (Fig. 2D). A trajectory analysis through the 500-ns simulation showed that Glu213, Phe214, and Pro215 underwent a marked conformation change after 340 ns (Fig. S4A). At 300 ns, the structure of the S2-S3 loop had essentially retained the initial crystal structure (Fig. S4B). In the 400-ns snapshot, by contrast, the location of the sidechain of Phe214 had largely shifted to the interior of the H2 helix, a hydrophobic environment made by Met211 (Fig. S4C). Pro215 had shifted to the vicinity of the D-ring, and the sidechain of Glu213 had swung around. We also investigated the dynamic character of the clustered waters over a very short timescale. Inspection of the trajectory showed that the water molecules forming a hydrogen bond with Glu217 (WAT21 and WAT74, in Fig. S5), and one of the clustered waters (WAT105, in Fig. S5) rapidly moved to the outside of the protein within 1 ns (Fig. S5). These analyses demonstrated the structural flexibility of the S2-S3 loop, and the mobility of the water cluster.

### Determination of the bilin protonation state by NMR

To evaluate the chemical environment of the PCB molecule, especially its protonation state, we performed NMR experiments using isotopically labeled PCB incorporated into the unlabeled GAF domain. First, we tried to detect NH signals of the pyrrole rings by 2D ^1^H-^15^N HSQC experiments. However, we observed only one signal from a pyrrole ring, probably assigned to N_D_-H, in both the Pr state and the Pg state: namely, (^1^H,^15^N) = (6.71, 134.4 ppm) and (^1^H,^15^N) = (8.82, 134.9 ppm), respectively, at 30 °C and pH 7.5 (Fig. S6). This contrasts with previous NMR studies of the red/green CBCR NpR6012g4 (21, 50), where four ^15^N-^1^H signals were observed in both the Pg and the Pr state in 2D ^1^H-^15^N HSQC experiments. These results imply that the exchange rates of the NH moieties NA, NB, and NC of PCB are much faster in RcaE than in NpR6012g4.

We therefore tried to detect ^15^N signals directly by a 1D ^15^N experiment, which does not need a magnetization transfer step from ^1^H to^15^N. Although direct detection of ^15^N is intrinsically insensitive, we overcame the low sensitivity by using a cryogenic broad-band observation probe. Via this strategy, we successfully observed four ^15^N signals (I, II, III, IV) within 10 hours at 30 °C in the Pr state, and four ^15^N signals (I’, II’, III’, IV’) in the Pg state (Fig. 3A and B). The chemical shifts of I, II, III, and IV were 134.9, 160.4, 163.5, and 163.5 ppm, respectively, with III and IV overlapping (see supporting text and Fig. S7), and those of I’, II’, III’, and IV’ were 134.4,152.5, 156.3, and 258.1 ppm, respectively. The signals at 130–160 ppm corresponded to a protonated Schiff base, while signal IV’ at 258.1 ppm in the Pg state alone showed a large low-field shift, corresponding to a deprotonated Schiff base. This indicates that one nitrogen atom of PCB is deprotonated in the Pg dark state, whereas all nitrogen atoms are protonated in the Pr photoproduct state. We also performed 1D ^15^N measurement of the Pr state of RcaE under conditions of pH 9 and pH 10, where RcaE is converted to the green-absorbing Pg state harboring C15*-E* PCB (18). Strikingly, the low-field shifted peak at ~258 ppm was clearly observed even in the C15*-E* Pg state at both pH 9.0 and pH 10 (Fig. 3C and D). This result supports the idea that the peak at 258.1 ppm in the Pg state at pH 7.5 is the signal from deprotonated PCB.

**Figure 3.**
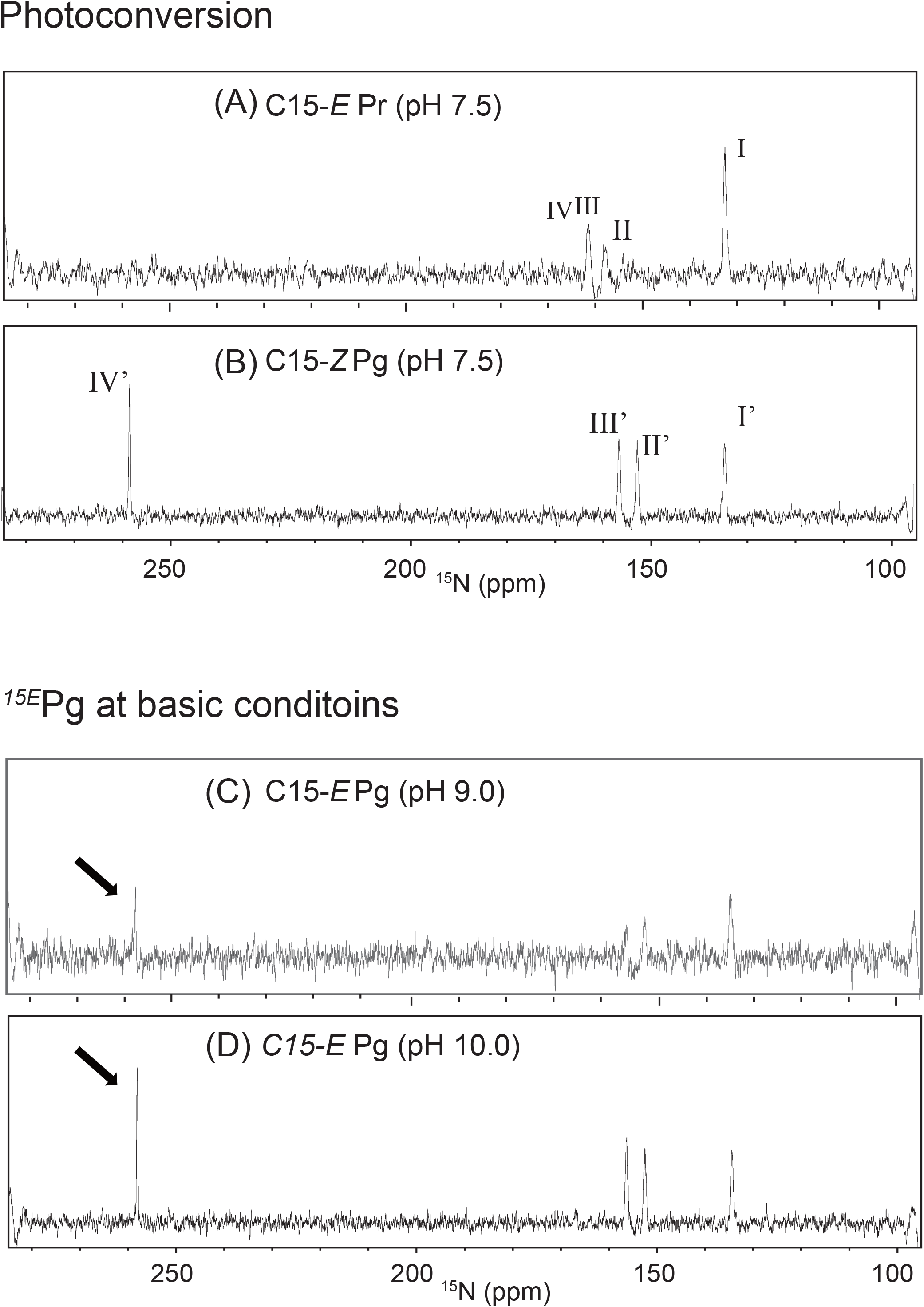
NMR spectra of the nitrogen atoms of PCB. (A)(B) Spectra recorded before and after photoconversion. Shown is the ^15^N 1D NMR spectrum of PCB in the Pr state (A) and the Pg state (B). (C)(D) pH titration in the Pr state. Shown is the ^15^N 1D NMR spectrum of PCB in the Pr state at pH 9.0 (C), and pH 10 (D).

The assignments of ^15^N signals are informative for understanding which nitrogen atom is protonated or deprotonated. However, the lack of an NH correlation makes it difficult to establish the assignments experimentally (Fig. S6). To obtain clues, we performed quantum mechanics/molecular mechanics (QM/MM) calculations to simulate the ^15^N NMR spectra (Fig. S8A-F). For the Pr state, the MD snapshot at 270 ns, which was closest to the averaged structure of the MD trajectory, was used to perform a subsequent geometry optimization. We computed the ^15^N chemical shifts by using the gauge-independent atomic orbital (GIAO) method (51, 52). The chemical shifts were also calculated for the Pg state. For the latter case, we used two reported Pg models in which C15-*Z*,*anti* PCB is deprotonated at N_B_ or N_C_ (39). As shown in Fig. S8, the ^15^N 1D spectra of both the Pr (Fig. S8A) and Pg (Fig. S8B) states were well reproduced by the Pr and N_B_-deprotonated Pg models, respectively. Notably, the chemical shift calculated with the N_C_-deprotonation (Fig. S8C) deviated from the corresponding experimental value compared to that of N_B_-deprotonation (Fig. S8B). Further, the calculation in which PCB is deprotonated at N_B_ or N_C_ based on the structure of C15-*E*,*syn* crystal structure was also conducted. We constructed initial structures for the deprotonated RcaE from the present crystal structure. The calculated specta (Fig. S8 D and E) deviated more from the observed spectra as compared with the calculated spectra for models in which in C15-*Z*,*anti* PCB (Fig. S8B and C).

On the basis of these results, signals I, II, III, and IV for Pr were assigned to N_D_, N_B_, N_A_, and N_C_, respectively (or N_D_, N_B_, N_C_, and N_A_, because III and IV overlapped). On the other hand, signals I’, II’, III’, and IV’ for Pg were assigned to N_D_, N_A_, N_C_, and N_B_, respectively, suggesting that deprotonation in the Pg state occurs at N_B_. These QM/MM calculations (Fig. S8A-E) were performed using an electronic embedding (EE) scheme, where electrostatic environments of the protein moiety are included in the QM calculations. To explore the effects of these electrostatic environments of the protein including hydrogen bonds, we next performed a QM/MM calculation of the Pr model with a mechanical embedding (ME) scheme, which does not incorporate the electrostatic interactions between the QM and MM regions. A distinct difference between the EE and ME schemes was observed for the calculated spectra (Figs. S8A and F), indicating that the ^15^N chemical shifts reflect not only the chromophore structure but also the electrostatic environment of the protein moiety.

## Discussion

We have reported a high-resolution (1.63 Å) crystal structure of the bilin-binding GAF domain of the chromatic acclimation sensor RcaE, a representative member of the green/ red CBCR subfamily. The PCB molecule adopts a unique C5-*Z,syn*, C10-*Z,syn*, C15*-E,syn* structure and is buried in the hydrophobic pocket of the GAF domain. We identified a direct hydrogen bond network connecting the NA–NC nitrogen atoms of PCB, the carboxyl group of Glu217, and the pentagon water cluster in a unique cavity within the PCB-binding pocket. MD simulation suggested the rapid exchange of the clustered waters with solvent and the flexibility of the S2-S3 loop. 1D 15N NMR analysis demonstrated that the four nitrogen atoms of PCB are indeed fully protonated in the Pr photoproduct state, and one of them is deprotonated in the Pg dark state. These findings uncover the structural basis of the protochromic green/red photocycle.

NMR is a powerful approach because it can provide direct information on the chemical environments of PCB, including its protonation state. We used 1D ^15^N measurements and demonstrated that one of the four NH moieties of PCB is actually deprotonated in the C15-*Z* Pg dark state created by illumination with red light and also in the C15-*E* Pg photoproduct state formed under basic pH conditions (Fig. 3). The similarity in the down-filed shift of ^15^N in the C15-*Z* Pg and C15-*E* Pg states suggests that the PCB chromophore undergoes common structural changes by deprotonation, which affects the electronic state of the π-conjugated system of PCB. Comparison of the observed chemical shift of ^15^N and the calculated chemical shifts using the quantum chemical calculations suggested that deprotonation occurs at the N_B_ nitrogen (Figs. 3 and S8), which is consistent with our previous resonance Raman spectroscopy and QM/MM simulation experiment in RcaE (53). Deprotonation in the N_B_ nitrogen is also consistent with solid-state NMR of *in vitro*-prepared AnPixJ proteins harboring deprotonated PCB fraction for its Pg photoproduct state (54). In 2D ^1^H-^15^N HSQC measurement, three of the four NH signals were missing for RcaE (Fig. S6), whereas all four NH signals were observed for the red/green CBCR NpR6012g4 (21, 50). This difference probably reflects the fast exchange of the proton in the NH moieties of the PCB molecule. Our structure suggested that the three N_A_-N_C_ pyrrole nitrogens form direct hydrogen bond with E217, whereas the N_D_ positioned rather hydrophobic environment (Figs. 1E and F). Therefore, we speculate that the rapid exchange of the proton occurs in the N_A_-N_C_ of the PCB. Ultrafast transient absorption spectroscopy suggested the presence of ground-state inhomogeneity in the Pg dark state of RcaE, which was previously interpreted as the existence of different tautomers: namely, N_B_- and N_C_-deprotonated PCB (55). Although we did not observe apparent heterogeneity of the Pg state in 1D ^15^N measurements (Fig. 3), this measurement is less sensitive in detection of the ground state heterogeneity due to its low S/N signals and changes of the intensity of each peak caused by the protonation state (Fig. 3), We also could not rule out the possibility that averaging of the signals between N_B_-deprotonation and N_C_-deprotonation.

Previously, we performed extensive site-directed mutagenesis of residues in the GAF domain of RcaE and investigated the effect of these substitutions on the bilin p*K*_a_ by monitoring of the pH-induced Pg/Pr conversion (18). Taking these results together, we elucidated the detailed roles of each residue in the PCB-binding pocket of RcaE. In the Pr structure, the carboxyl group of Glu217 forms direct hydrogen bonds with the N_A_-N_C_ pyrrole nitrogen atoms (Fig. 1E and F). The E217A and E217Q mutants showed the normal Pg formation but were deficient in Pr formation, whereby the upshift in bilin p*K*_a_ in the Pr state was specifically inhibited (18). By contrast, the E217D mutant of RcaE showed a near normal Pg/Pr conversion and p*K*_a_ shift (18). Therefore, the role of the carboxyl group of Glu217 is to stabilize the protonation of NH moieties with its negative charge. Although pyrrole nitrogens are hydrogen bonded with carboxylate in other CBCR subfamilies, the uniqueness of RcaE is the presence of the “holey pan” structure in the Pr photoproduct state identified herein (Fig. 2C). In this structure, the carboxyl group of Glu217 also forms a direct hydrogen bond with the clustered water, which is rapidly exchanging with solvent in the MD simulation (Fig. S5). In addition, the fast exchanging of the proton in the N_A_-N_C_ pyrrole nitrogen atoms was implied by the absence of the three of four NH signals in 2D ^1^H-^15^N HSQC measurement (Fig. S6). Therefore, we propose that the unique holey pan structure provides PCB with a route to access solvent waters, which supports the proton transfer and subsequent formation of the Pr photoproduct (Fig. 4, right panel). Notably, in this mechanism, proton can be supplied by the solvent water and a proton donor of a specific charged residue is not required.

**Figure 4.**
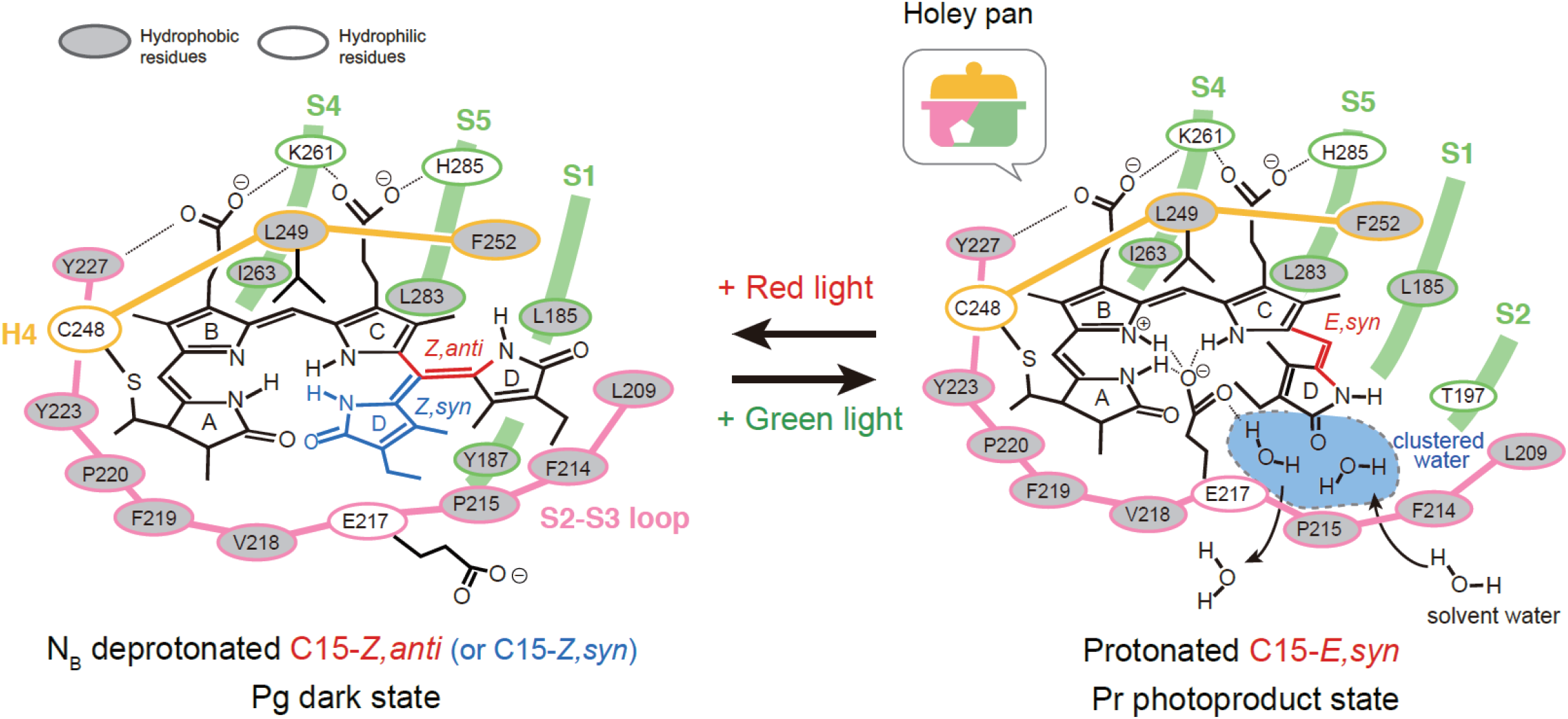
Proposed model of the photoconversion of PCB in RcaE. In the C15*-E* Pr state, PCB adopts a *syn* conformation and all nitrogen atoms are protonated. In the C15*-Z* Pg state, PCB adopts an *anti* conformation and N_B_ is deprotonated while the other nitrogen atoms are protonated.

Upon red light illumination, RcaE undergoes C15-*E* to C15-*Z* photoisomerization and subsequent deprotonation of PCB (downshift of the bilin p*K*_a_), resulting in formation of the Pg state (18, 38). The E217A and E217Q mutants showed normal p*K*_a_ value in Pg dark state (18). Therefore, it is less likely that the negatively charged carboxyl group functions as a base and decreases the bilin p*K*_a_ in the Pg state. In the Pg state, the carboxyl group of Glu217 may apart from PCB or be exposed on the surface of the protein. Notably, our structure shows that Leu249 is positioned toward the α-face of PCB and forms a hydrophobic lid, together with Phe252 (Figs. 1E and F). The L249H mutant showed a substantial increase in bilin p*K*_a_ and formed the Pr state even in the C15-*Z* state, whereas F252H was not deficient in the Pg formation (18). These data suggest that the hydrophobicity of Leu249 is crucial for the deprotonation of PCB in the Pg state. It is known that the p*K*_a_ values of charged moieties shift in the direction that favors the neutral state in the hydrophobic protein core (56, 57). Therefore, we propose that a “desolvation effect” is the driving force behind the deprotonation of PCB. Lys261 forms ion pairs with both the B- and C-ring carboxylates, together with Tyr227 and His285 (Figs. 1E and F). Previously, the Y227F and H285Q mutants showed negligible effects on p*K*_a_ regulation, whereas both K261A and K261M showed substantially decreased p*K*_a_ in both the Pg and Pr states (18). The presence of the hydrophobic lid of Leu249 and Phe252 in our structure suggests that Lys261 is not likely to function as a proton donor as discussed previously (18), but rather tunes the bilin p*K*_a_ by adjusting the location of the C15-*E,syn* PCB molecule in the GAF domain (Fig. 4).

For the PCB structure in the Pg dark state, whose three-dimensional structure is not yet available, we can argue two possible structures: C15-*Z,anti* and C15-*Z,syn* (Fig. 4). The C15-*Z,anti* structure for the Pg state of RcaE (Fig. 4, left panel) is supported by recent our resonance Raman spectroscopy of RcaE (53). That study showed that the observed resonance Raman spectra in the Pg state of RcaE were well reproduced by the simulated spectra of C15-*Z,anti* PCB in the N_B_-deprotonated state using a QM/MM approach combined with MD simulation (53). The C15-*Z,anti* structure is also supported by the recent *in virtro* reconstitution experiment (18). This study showed that apo-RcaE can incorporate a synthetic PCB analog that is sterically locked in the C15-*Z,anti* structure and forms a photoinert and red-shifted Pg state (18). The structure of C15-*Z,anti* PCB differs from natural PCB in that (1) it has a bulky D-ring with an additional covalent bridge between C13 of the C-ring and N_D_ of the D-ring, and (2) its C-D ring forms a nearly planar structure and is unable to undergo structural changes owing to the presence of dual covalent linkages (58). Incorporation of C15-*Z,anti* PCB into the apo-protein of our Pr structure would cause a steric clash with Leu209 and Phe214 in the S2-S3 loop (Fig. S2). Therefore, the acceptance of the C15-*Z,anti* PCB suggests the occurrence of a structural rearrangement of the S2-S3 loop in the Pg dark state. The flexibility of the S2-S3 loop and the presence of the interior cavity inside the S2-S3 loop containing a water cluster may enable the structural change of PCB along with C15-*E,syn* to C15-*Z,anti* (probably through C15-*Z,syn* intermediate).

By contrast, the C15-*Z,syn* model for the Pg state of RcaE (Fig. 4, left panel) is consistent with the recent crystal structure of the Pfr dark state of 2551g3 of a far-red/orange subfamily, where the PCB bilin adopts a C5*-Z,syn*, C10*-Z,syn*, C15*-Z,syn* structure (25). Phylogenetic analysis showed that the far-red/orange subfamily evolved from the green/red subfamily (59). RcaE shares most of residues in the PCB-binding pocket with 2551g3 including Leu249 and Phe252 forming a hydrophobic lid, Tyr227, Arg233, Lys261, and H285 interacting with B- and C-ring carboxylates, and Glu217 interacting with the N_A_-N_C_ pyrrole nitrogens (Fig. S2). In contrast, residues on the S2-S3 loop are divergent between RcaE and 2551g3 except for Glu217 (Fig. S2), and there is no cavity in the Pfr state of 2551g3 (25). Therefore, the variation of the residues in the S2-S3 loop must be responsible for their distinct spectral sensitivity between the green and far-red region, corresponding to a 200-nm difference in absorption. pH titration showed that the C15*-Z,syn* PCB is protonated in the Pfr state of 2551g3, where Glu914 (corresponding to Glu217 in RcaE) forms hydrogen bonds with the four pyrrole nitrogens (N_A_-N_D_). Site-directed mutagenesis demonstrated that Glu914 is required for the protonation in the Pfr state (25). From these results, Bandara et al proposed that the far-red shift of the Pfr dark state of 2551g3 has a di-protonated lactim isomer of the A-ring with the deprotonated N_B_ nitrogen (25). This model has not yet been investigated by vibrational and NMR spectroscopies, but our 1D ^15^N NMR data agree with the N_B_ deprotonation model in 2551g3. If the RcaE has a C15-*Z*,*syn* structure for the Pg dark state as shown in the 2551g3, PCB undergoes structural changes between C15*-Z,syn* and C15*-E,syn* upon photoconversion, which requires less drastic structural changes compared with the C15*-Z,anti* model.

In summary, we propose a revised model of the protochromic green/red photocycle of RcaE (Fig. 4). In the Pr state, PCB adopts a C15-*E,syn* structure and the N_A_-N_C_ pyrrole nitrogen atoms form a direct hydrogen bond with the carboxyl group of Glu217. The carboxyl group of Glu217 also forms a direct hydrogen bond with solvent water molecules in the holey pan structure, which increases hydrophilicity of the vicinity of PCB and leads to the protonation as well as Pr formation. Red light illumination causes a structural change of PCB from C15-*E,syn* to C15-*Z,anti* (or C15-*Z,syn*) with release of the carboxyl group of Glu217 from PCB. The accessibility of PCB with solvent decreased by the disconnection of the direct hydrogen bond with E217 and the presence of hydrophobic L249 in a-face, which lead to the decrease of the bilin p*K*_a_. Deprotonation of the N_B_ of PCB causes a change in the electrostatic state of the π-conjugated system of PCB, resulting in the drastic absorption change from the red to the green region. For reverse reaction, Green light illumination causes C15-*Z,anti* (or C15-*Z,syn*) to C15-*E,syn* conversion and protonation of the N_B_. In this model, the carboxyl sidechain of the Glu217 functions as a gate in a proton-exit and proton-influx pathway to solvent water. We would favor the C15-*Z,anti* model that is supported by the spectroscopic evidence, but three-dimensional structure determination of RcaE in the deprotonated Pg dark state is required to settle the discussion of the D-ring structure.

## Materials and Methods

### Sample preparation

We prepared the GAF domain as previously descried (18). In brief, the GAF domain (residues 164–313) of RcaE was cloned into plasmid pET-28 (Novagen). The construct was expressed in *Escherichia coli* strain BL21(DE3) star (Invitrogen) as a fusion protein with a His tag, together with PCB biosynthetic plasmid pKT271 (41). The harvested cells were lysed via sonication. The His-tag protein was purified by using a column of Ni-Sepharose 4B (Qiagen), followed by gel filtration using a Superdex 75 column (G.E. Healthcare) equilibrated with a 10 mM Tris-HCl (pH 7.5) buffer containing 50 mM KCl. For crystallization, the protein was further purified by using a HiTrap-SP ion-exchange column (G.E. Healthcare), equilibrated with a 20 mM Tris-HCl (pH 8.0) buffer containing 50 mM KCl and eluted with a gradient of 1M KCl. The purified sample was stored in buffer containing 20 mM Tris-HCl (pH 7.5) and 10 mM NaCl.

### Crystallization

Prior to crystallization experiments, the protein solution of RcaE was irradiated with green LED light to adapt it to the Pr photoproduct state. Crystallization experiments were carried out under green light, and the crystallization plates were incubated in a dark environment. Crystals of the red-absorbing form were obtained at 293 K via the hanging-drop vapor diffusion method with drops containing of 0.9 μL of protein solution (25 mg/mL in 20 mM TrisꞏHCl, pH 7.5, 10 mM NaCl) mixed with 0.9 μL of reservoir solution (100 mM TrisꞏHCl, pH 7.2, 200 mM MgCl2, 27% PEG 4000). The blue plate-shaped crystals grew to typical dimensions of 200 × 50 × 20 μm3 within 1 week.

### Data Collection and Structure Refinement

Data collection was performed at 95 K under green LED light on Nagoya University beamline BL2S1 at the Aichi Synchrotron Radiation Center (AichiSR), Japan (60). The diffraction pattern was indexed, integrated, and scaled by using XDS (61). We confirmed the blue color of the crystal after X-ray irradiation (Fig S9). The initial structure was solved by MOLREP (62) as implemented in CCP4 (63) using the TePixJ GAF domain (PDB: 3VV4) as a search model.

Refinement and manual correction of the structure were performed by using REFMAC5 (64) and Coot (65), respectively. For analyses of the PCB configurations and the clustered waters, simulated-annealing refinement in PHENIX were performed. Parameters for data collection, processing, and structure refinement are listed in Table S1. Figures were drawn by using PyMOL (the PyMOL Molecular Graphics System, Schrödinger, LLC).

### Data Availability

The X-ray diffraction dataset and associated model have been deposited in the Research Collaboratory for Structural Bioinformatics (RCSB) Protein Data Bank (https://www.rcsb.org) under PDB ID code 7CKV.

### NMR

Uniform labeling of PCB with 15N and 13C was achieved by using M9 minimal medium containing no stable isotope and uniformly labeled 15N and 13C 5-aminolevulinic acid. Prior to plasmid induction, the isotope-enriched 5-aminolevulinic acid was added to M9 medium. The purified sample was dissolved in 50 mM Tris-HCl buffer (pH 7.5) containing 50 mM KCl in 95% H2O/5% 2H2O. For pH titration, buffers containing 50 mM CHES (pH 9.0) plus 50 mM KCl, and 50 mM CHES (pH 10.0) plus 50 mM KCl were used. The final concentration of the protein was approximately 1 mM. NMR spectra were acquired at 30°C on a Bruker AVANCEIIIHD 500 instrument equipped with a cryogenic BBFO probe, and on a Bruker AVANCEIIIHD 600 instrument equipped with a cryogenic TCI probe. The 15N 1D experiment was achieved with 1H decoupling during acquisition. 2D 1H-15N HSQC spectra were acquired by using a standard gradient-echo sensitivity improvement pulse sequence with a WATERGATE-Water-flip-back type water suppression technique. Data processing was performed with Topspin ver. 3.5 (Bruker).

### MD Simulation

Both the initial setup and the MD runs were performed with the Amber16 program (66) using an explicit representation of solvent molecules and the ff14SB all-atom force field under periodic boundary conditions. The initial starting geometry was taken from the crystal structure of the GAF domain of RcaE in the Pr state. The whole system was subjected to minimization and heated to 300 K. The system obtained after heating was simulated for 500 ns at 1 atm.

### QM/MM Calculation

The QM/MM calculations were performed by using Gaussian16 (67). The QM region consists of the PCB chromophore, while the remainder of the system was treated as the MM region. The initial geometry was obtained from a snapshot of the MD simulation, and 100 water molecules near the surface of the protein were also included. The QM part of the system was computed at the B3LYP/6-31G* (geometry optimization) or Hartree-Fock/6-311þG(2d,p) (chemical shifts) level of theory. The MM part was described by the Amber force field (68), and the positions of the MM atoms were frozen during geometry optimization. In most cases, an EE scheme that considers the partial charges of the MM region into the QM Hamiltonian was used. The GIAO nuclear magnetic shielding values were calculated, and the chemical shifts for the nitrogen atoms were obtained by subtracting their chemical shielding values from that calculated for NH_3_.

## Supporting information

Supplementary test and Tables

## Acknowledgments

We thank Dr. J. Clark Lagarias and Dr. Nathan C. Rockwell for helpful discussions. We thank Dr. Masatsune Kainosho for helpful discussion on the ^15^N 1D NMR experiment. This work was supported by JSPS KAKENHI (Grant Numbers 19H05645 and 19H05773 to Y.I., 19K06707 to Y.H. and 19H03169 to M.M.). The crystallography experiment at BL2S1 was performed under the approval of Aichi Synchrotron Radiation Center (Proposal No.2019N5012), and the NMR experiments were performed in part using the NMR spectrometers with the ultra-high magnetic fields under the Collaborative Research Program of Institute for Protein Research, Osaka University, NMRCR-18-05, −19-05 and −20-05. Some of the computations were performed at the Research Center for Computational Science, Okazaki, Japan.

