## Supplementary test and Tables for "Structural basis of the protochromic green/red photocycle of the chromatic acclimation sensor RcaE"

[1]Masaki Mishima

[1]Yuu Hirose

[1]Masashi Unno

Takayuki Nagae: 0000-0001-7016-5183

Tomotsumi Fujisawa: 0000-0002-3282-6814

Masashi Unno: 0000-0002-5016-6274

Yohei Miyanoiri: 0000-0001-6889-5160

Yuu Hirose: 0000-0003-1116-8979

Kei Wada: 0000-0001-7631-4151

Yutaka Ito: 0000-0002-1030-4660

Masaki Mishima: 0000-0001-7626-7287

**Classification**

Biological Sciences

**Keywords**

cyanobacteriochrome, phytochrome, bilin, NMR, crystallography

### Supporting text

#### Overlapping $^{15}N$ 1D signals in the Pr state

The ratio of the integration values of signal II and the overlapping signals III–IV was 1.0 to 1.9 (Fig. S7), indicating that signal II was derived from one nitrogen atom, while signals III–IV were derived from two nitrogen atoms. Of note, the integration value of signal I was 2.7, which probably reflects a stable NH bond. This is consistent with the corresponding signal observed in 2D $^{1}H$-$^{15}N$ HSQC.  The stable bond should be shorter than labile bonds, effectively causing T1 relaxation of the nitrogen atom.

**Supplemental figures and tables**

**Figure S1. UV-vis absorption spectrum of the GAF domain.** Shown is the UV-vis absorption spectrum of the $^{15Z}$Pg (green line) and $^{15E}$Pr (red line) states of the GAF domain from RcaE at pH 7.5.

**Figure S2. Sequence alignment of the GAF domain from green/red type and far-red/X type CBCRs.** The secondary structure elements of RcaE are indicated above the alignment. Hydrophobic (yellow) and hydrophilic (red) residues forming the PCB-binding pocket are shown. Residues conserved specifically in the S2-S3 loop (blue box) of the green/red subfamily are shown (green). PCB-anchoring Cys residue (star) and protochromic triad maintaining the bilin pKa (triangle) are shown below the alignment. Protein orthologs and host organisms are as follows. FdRcaE from *Tolypothrix* sp. PCC 7601/1 (*Fremyella diplosiphon*), Sc7335_RcaE from *Synechococcus* sp. PCC 7335, Lepjsc1_RcaE from *Leptolyngbya* sp. JSC-1, SyCcaS from *Synechocystis* sp. PCC 6803, NpCcaS from *Nostoc punctiforme* ATCC 29133, Lep6406_CcaS from *Leptolyngbya* sp. PCC 6406, Gm3708_CcaS from *Geminocystis* sp. NIES-3708, Gm3709_CcaS from *Geminocystis* sp. NIES-3709, Anacy_4718g3 and Anacy_2551g3 from *Anabaena cylindrica* (strain ATCC 27899 / PCC 7122), Cyan7822_4053g2 from *Cyanothece* sp. PCC 7822, Sta7437_1656 from *Stanieria cyanosphaera* PCC 7437, WP_016871037 from *Fischerella thermalis*, Syn7502_01757 from *Synechococcus* sp. PCC7502, and WP_046814686 from *Calothrix* sp. 336/3.

**Figure S3. MD trajectory analysis of ion pairs between PCB and the RcaE protein.** (A) Distance between the oxygen atoms (O1 and O2) of the B-ring propionate and the nitrogen atom of the Lys261 sidechain derived from the MD trajectory. (B) Distance between the oxygen atoms (OE1 and OE2) of the carboxylate of Glu217 and nitrogen atom $N_B$ derived from the MD trajectory.

**Figure S4. MD trajectory analysis of the S2-S3 loop.** (A) Root mean square deviation (RMSd) of heavy atoms in the MD snapshot structures versus the initial optimized structure. The RMSd for the total peptide backbone, the backbone without residues 270–275 (missing residues in the crystal structure), PCB, and the tripeptide region Glu213-Phe214-Pro215 (corresponding to the middle part of the S2-S3 loop_ is shown in black, blue, red and green, respectively. (B)(C) Ribbon models of the MD snapshot of Glu213-Phe214-Pro215 at 300 ns (B) and at 400 ns (C).

**Figure S5. MD trajectory of the clustered water.** (A) Ribbon and stick model of the initial optimized structure. (B) Ribbon and stick model of the MD snapshot at 1 ns. Asterisk indicates a water molecule penetrating into the cavity. (C) Distance between the oxygen atom of the carboxylate of Glu217 and either an oxygen atom of WAT21 derived from the MD trajectory within 1.5 ns (red) or an oxygen atom of WAT74 derived from the MD trajectory within 1.5 ns (blue); and distance between the backbone nitrogen atom of Phe214 and an oxygen atom of WAT105 derived from the MD trajectory within 1.5 ns (black).

**Figure S6. $^{1}$H-$^{15}$N 2D HSQC spectra of the GAF domain with selectively labeled PCB.** (A) $^{1}$H-$^{15}$N 2D HSQC spectrum of the PCB-labeled GAF domain in the Pr state. (B) $^{1}$H-$^{15}$N 2D HSQC spectrum of the PCB-labeled GAF domain in the Pg state. Arrow indicates the observed signals.

**Figure S7. 1D $^{15}$N spectrum of the GAF domain with selectively labeled PCB.** 1D $^{15}$N NMR spectrum of the PCB-selectively labeled GAF domain (repetition delay was set to ~6 sec). $^{15}$N signals (I, II, III, IV) and their integration values (colored in red) are indicated.

**Figure S8. Quantum mechanical calculation of $^{15}$N chemical shifts of PCB in the GAF domain.** Shown are the QM/MM $^{15}$N NMR chemical shifts calculated by using the gauge-independent atomic orbital**.** (A) Simulated spectrum of the C15-*E* Pr state of RcaE, in which all nitrogen atoms are protonated. (B) Simulated spectrum of the C15-*Z* Pg state of modeled RcaE, in which $N_B$ is deprotonated and $N_A$, $N_C$, and $N_D$ are protonated. (C) Simulated spectrum of the C15-*Z* Pg state of RcaE, in which $N_C$ is deprotonated and $N_A$, $N_B$, and $N_D$ are protonated. (D) Simulated spectrum of the C15-*E* Pg state of RcaE, in which $N_B$ is deprotonated and $N_A$, $N_C$, and $N_D$ are protonated. (E) Simulated spectrum of the C15-*E* Pg state of RcaE, in which $N_C$ is deprotonated and $N_A$, $N_B$, and $N_D$ are protonated. (F) Simulated spectrum of the C15-*Z* Pr state of RcaE, in which all nitrogen atoms are protonated with the mechanical embedding (ME) method.

**Figure S9. Pr crystal used for X-ray crystallographic analysis.** The photograph of the Pr crystal of the RcaE GAF domain was obtained after X-ray measurement. Scale bar corresponds to 0.1 mm.

**Movie S1. A video of 500 ns MD simulation of the Pr state of RcaE.** The PCB chromophore and the selected amino acid residues Glu217, Leu249, Phe252, Lys261, and His285 are illustrated by a stick model.

**Table S1.** Data collection and refinement statistics. Values in parentheses are for the outer shell.

| | |
|---|---|
| PDB Accession Code | 7CKV |
| Data collection | |
| Synchrotron, Beamline | AichiSR, BL2S1 |
| Temperature (K) | 95 |
| Space group | $P2_1$ |
| Cell dimensions (Å/°) | $a$ = 35.73, $b$ = 101.33, $c$ = 41.81, $\beta$=91.79 |
| Wavelength (Å) | 1.12 |
| Resolution (Å) | 38.66 - 1.63 (1.66 – 1.63) |
| Completeness (%) | 99.6 (99.9) |
| Observed reflections | 144213 (6993) |
| Unique reflections | 36876 (1810) |
| Redundancy | 3.9 (3.9) |
| Wilson B factor (Å$^2$) | 15.0 |
| $R_{merge}$ (%) | 4.6 (48.3) |
| CC1/2 (%) | 99.9 (83.7) |
| I/σI | 13.3 (2.2) |
| Refinement | |
| Resolution (Å) | 38.66 – 1.63 (1.67-1.63) |
| No. of reflections | 35093 (2589) |
| $R_{wiork}$ (%) | 15.8 (24.6) |
| $R_{free}$ (%) | 19.9 (22.7) |
| Average B factor (Å$^2$) | 20.2 |
| No. of atoms | 2754 |
| R.m.s. deviations | |
| Bond lengths (Å) | 0.011 |
| Bond angles (°) | 1.719 |
| Ramachandran statistics (%) | |
| Favored regions (%) | 99.6 |
| Allowed regions (%) | 0 |
| *Disallowed regions (%) | 0.4 |

*The Ramachandran outlier is Leu312 (C-terminal part) of chain B

**Table S2** Distances between the carbon atoms of PCB in the RcaE crystal structure, and the corresponding carbon atoms of TePixJ ($^{15E}$Pg) and Slr1393 ($^{15E}$Pg). The ring to which the carbon atom belongs (A, B, C or D) is given in parentheses. For reference, nomenclature (numbering of the carbon atom) of the PCB is shown at the bottom of the table.

| | C1 (A) | C4 (A) | C6 (B) | C9 (B) | C11 (C) | C14 (C) | C16 (D) | C19(D) |
|---|---|---|---|---|---|---|---|---|
| TePiIxJ | 3.8 | 3.2 | 4.3 | 4.4 | 4.3 | 3.9 | 3.1 | 4.6 |
| Slr1393 | 3.8 | 4.4 | 5.1 | 5.0 | 4.6 | 4.1 | 3.2 | 4.5 |

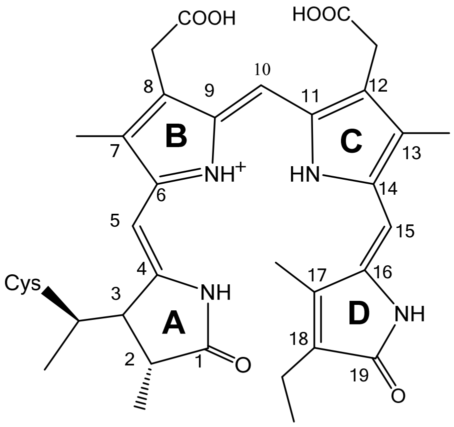
