## Supplementary figures and images for "Structural basis of the protochromic green/red photocycle of the chromatic acclimation sensor RcaE"

### Supplementary Figures

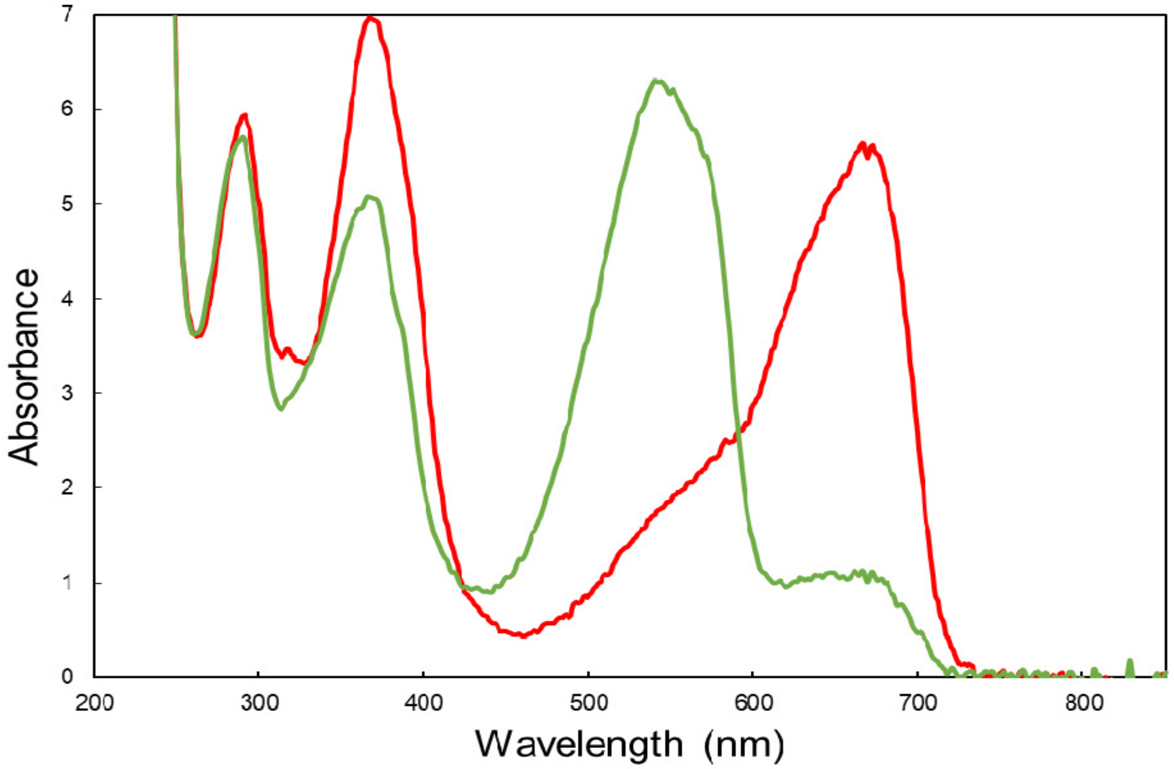

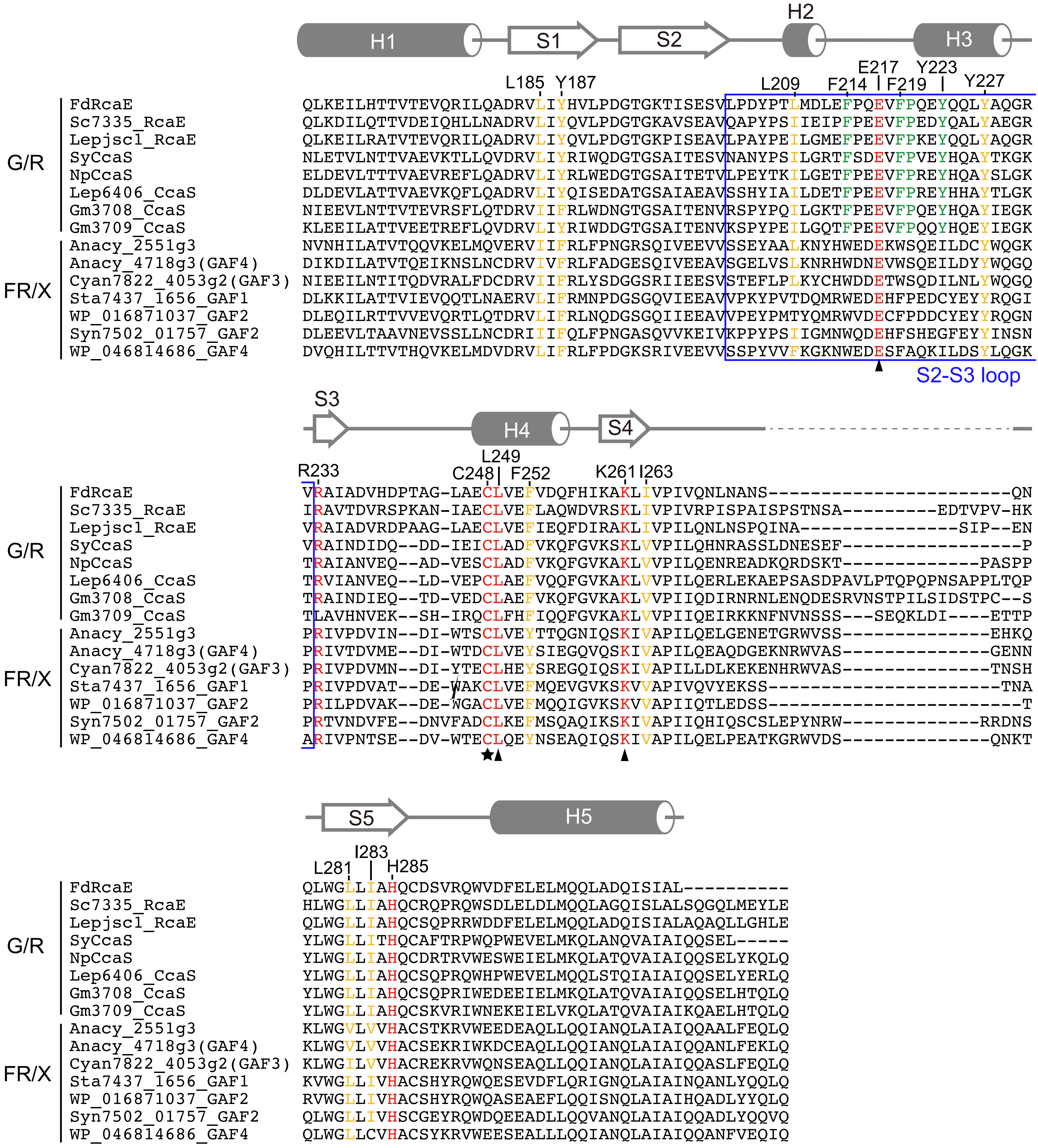

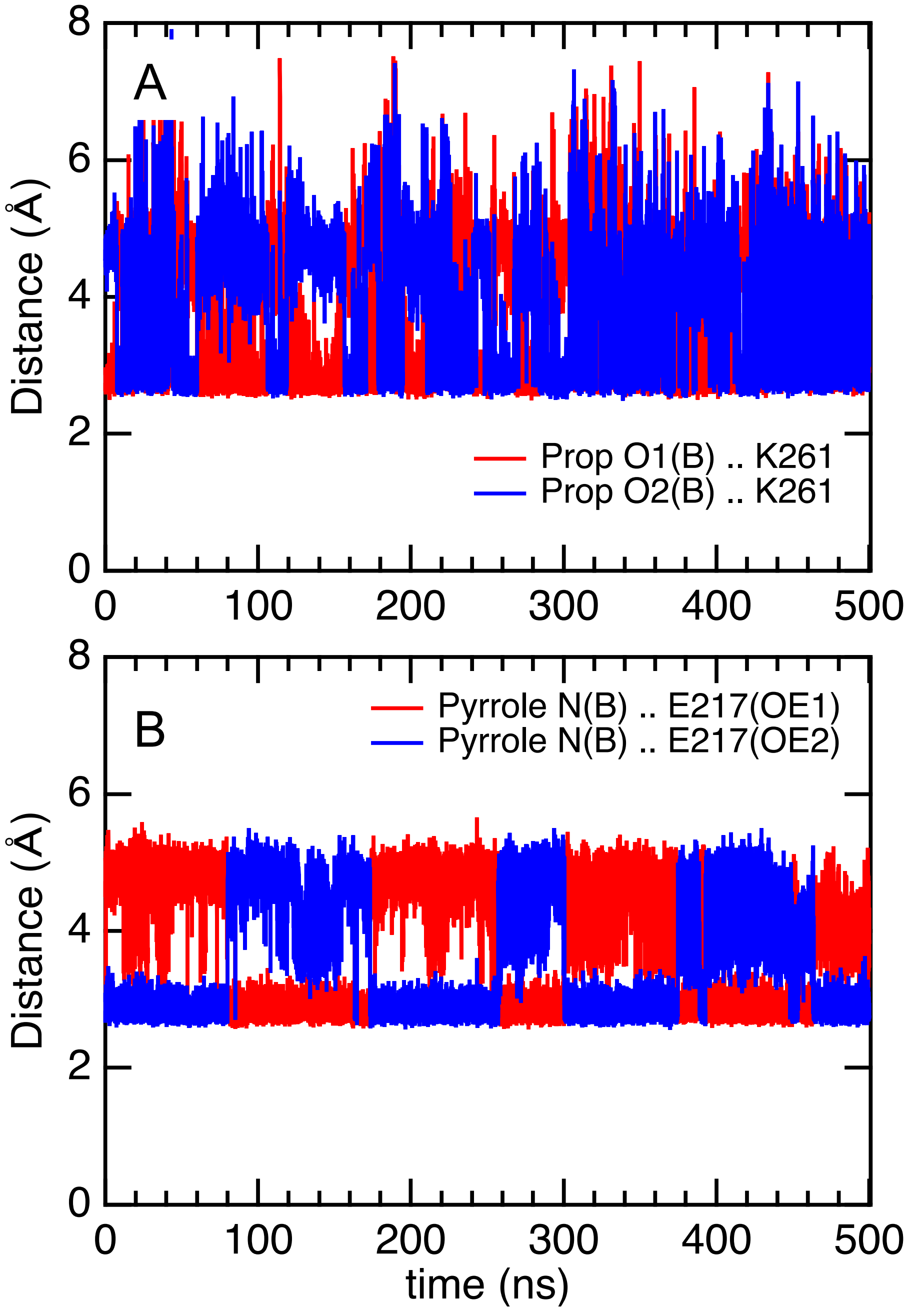

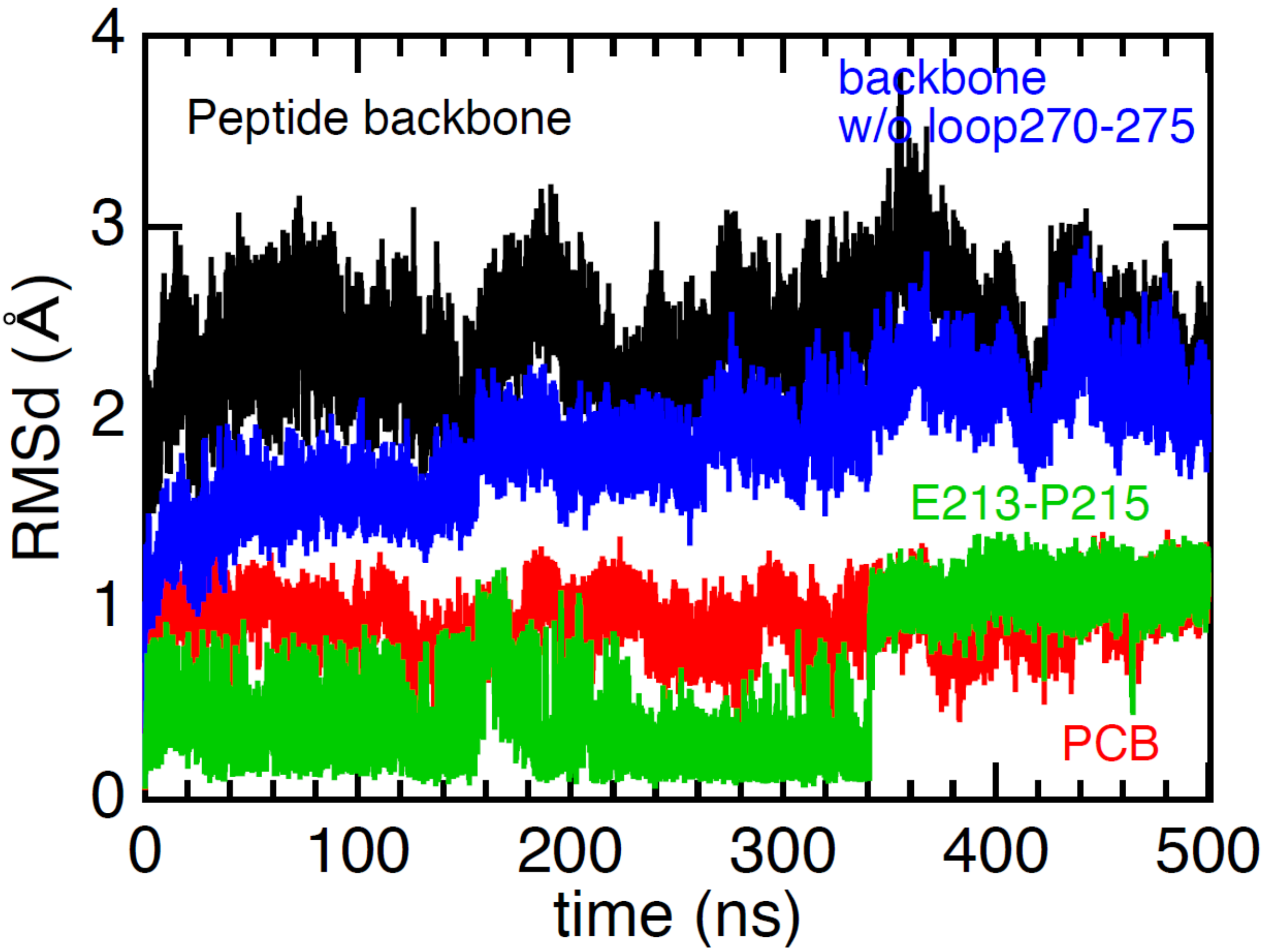


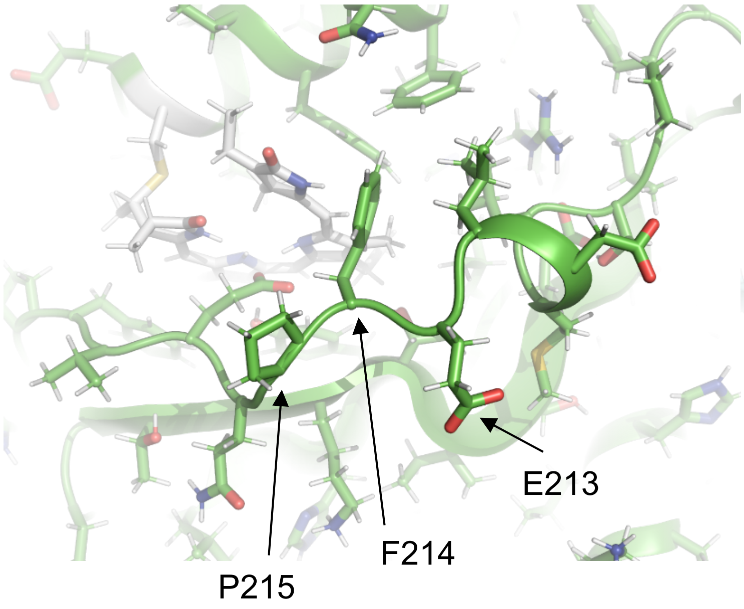


MD snapshot at 300 ns

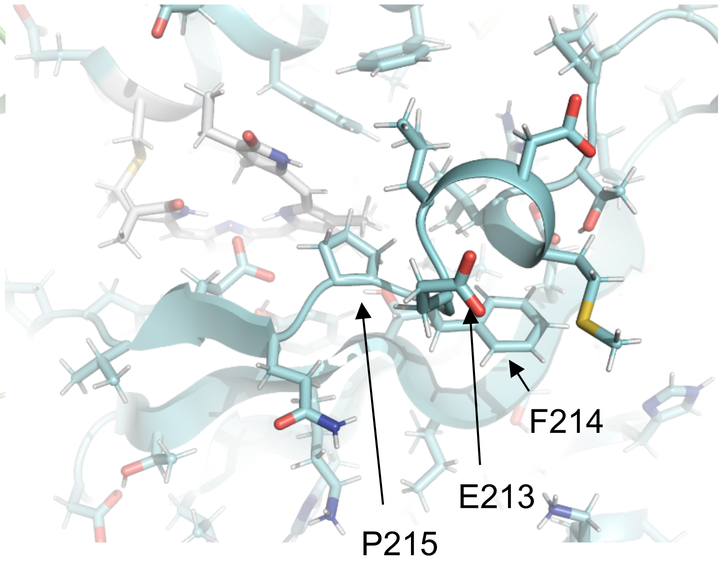


MD snapshot at 400 ns

Fig. S4

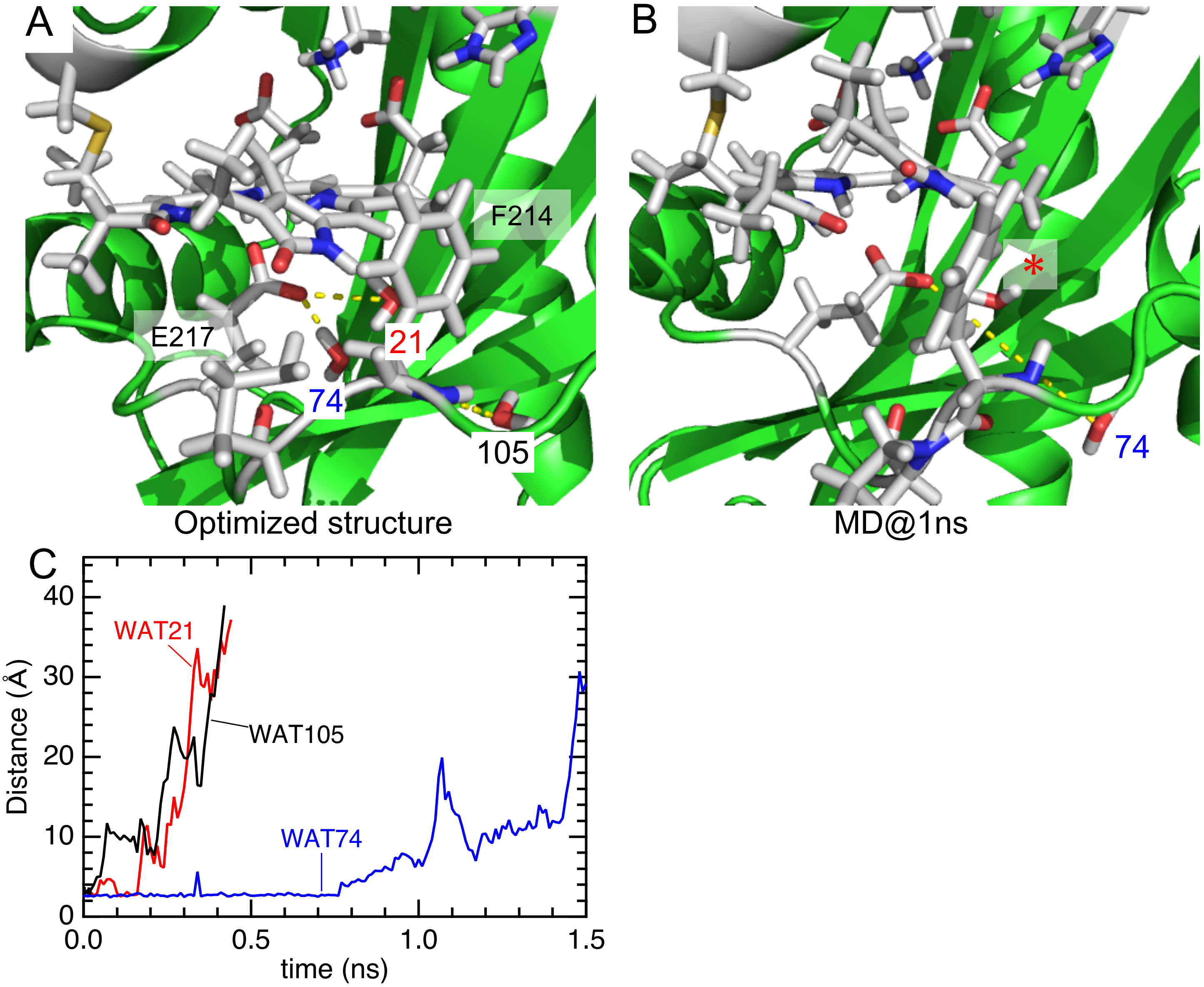

(A) $^{1}H$-$^{15}N$ 2D NMR (Pr)

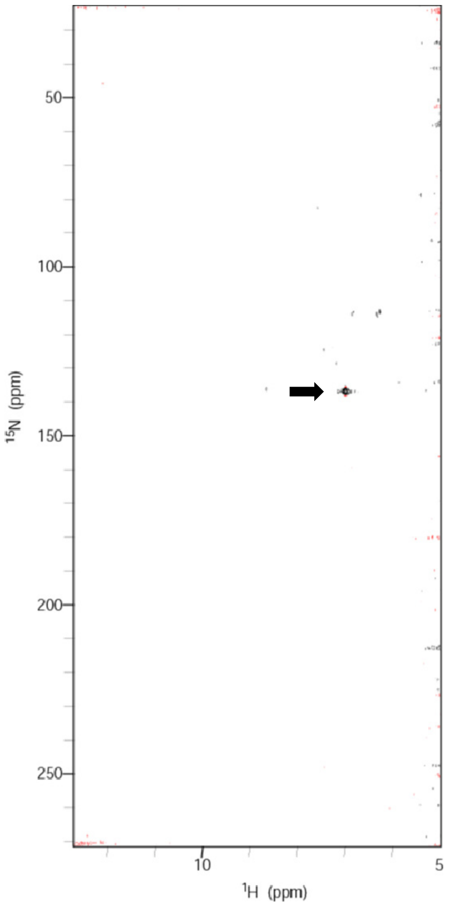


(B) $^{1}H$-$^{15}N$ 2D NMR (Pg)

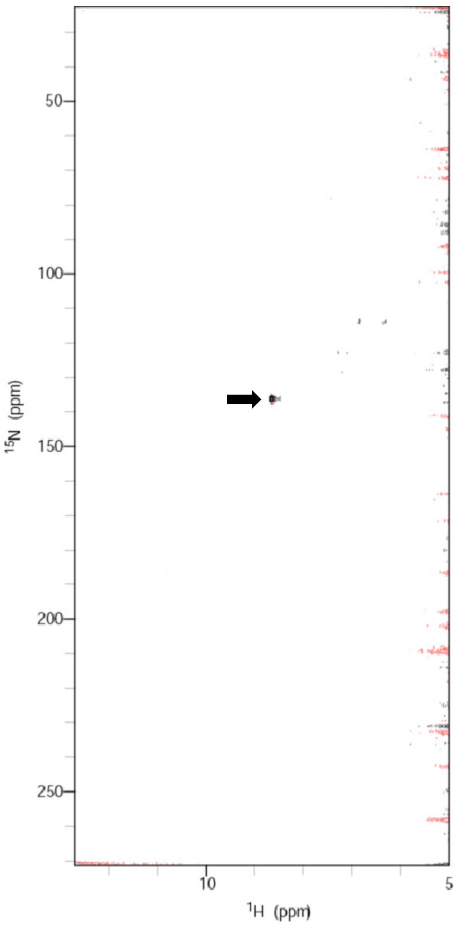


Fig. S6

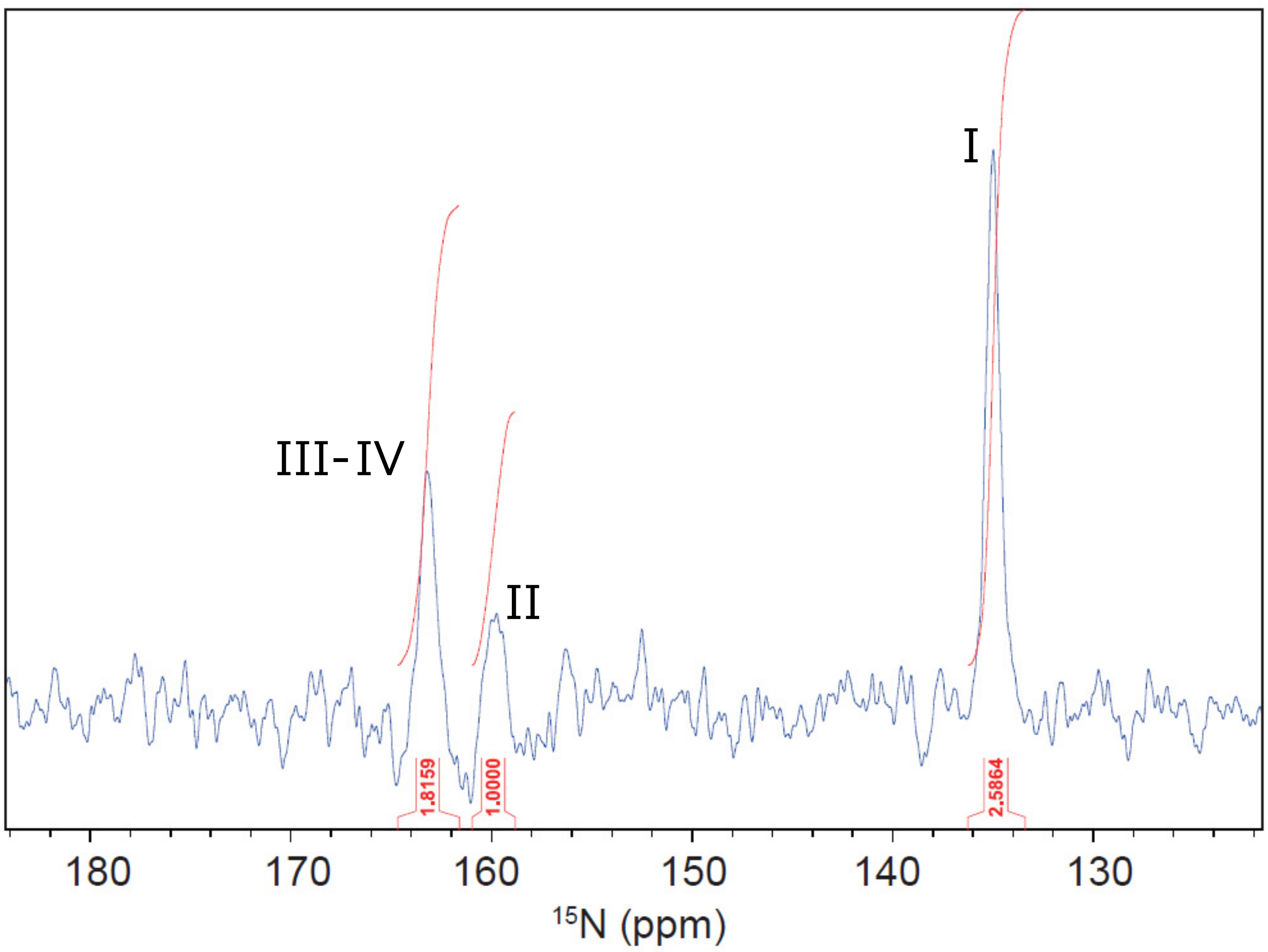


Fig. S7

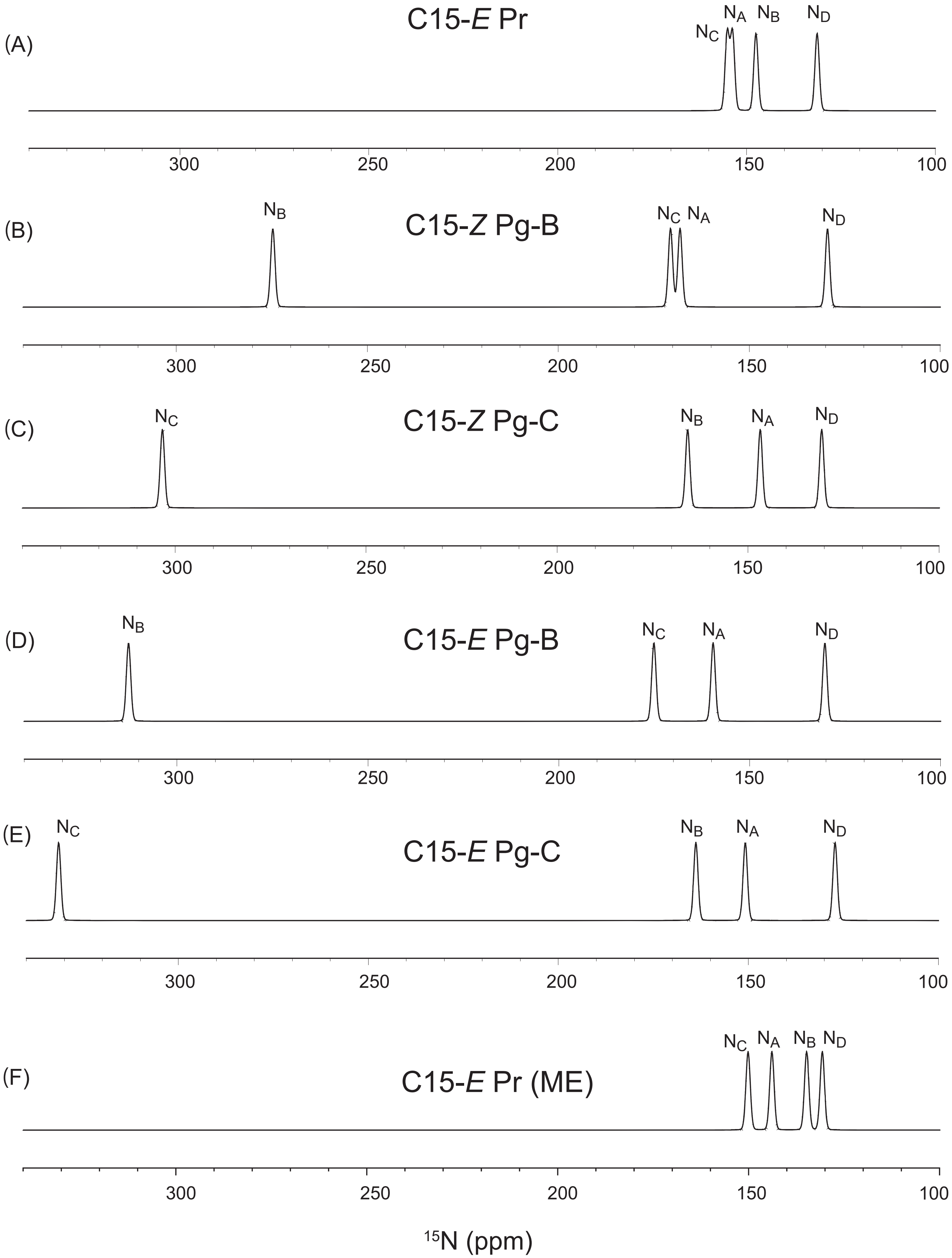

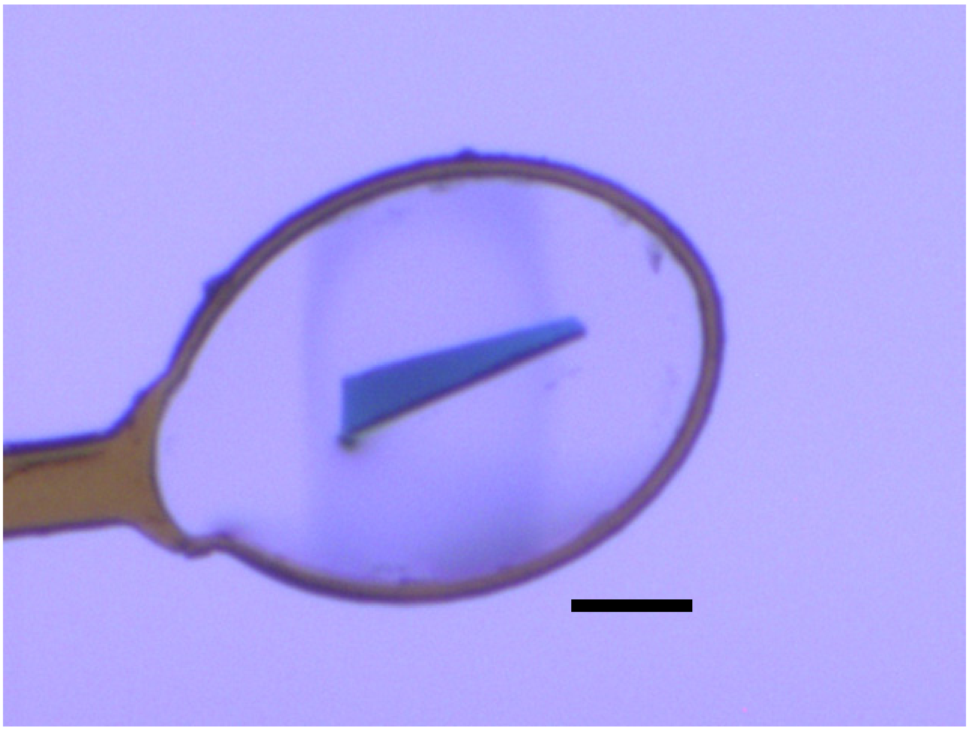

Fig. S9
